# Pharmacological determination of the fractional block of Nav channels required to impair neuronal excitability and *ex vivo* seizures

**DOI:** 10.1101/2022.06.08.494063

**Authors:** Samrat Thouta, Matthew G. Waldbrook, Sophia Lin, Arjun Mahadevan, Jannette Mezeyova, Maegan Soriano, Samuel J. Goodchild, R. Ryley Parrish

**Affiliations:** Department of Cellular and Molecular Biology, Xenon Pharmaceuticals, Burnaby, BC, Canada

**Keywords:** Sodium channel, inhibition, Epileptiform activity, tetrodotoxin, pharmacology, MEA

## Abstract

Voltage-gated sodium channels (Nav) are essential for the initiation and propagation of action potentials in neurons. Of the different channel subtypes, Nav1.1, Nav1.2 and Nav1.6 are prominently expressed in the adult central nervous system (CNS). All three of these sodium channel subtypes are sensitive to block by the neurotoxin tetrodotoxin (TTX), with TTX being almost equipotent on all three subtypes. In the present study we have used TTX to determine the fractional block of Nav channels required to impair action potential firing in pyramidal neurons and reduce network seizure-like activity.

Using automated patch-clamp electrophysiology, we first determined the IC_50_s of TTX on mouse Nav1.1, Nav1.2 and Nav1.6 channels expressed in HEK cells, demonstrating this to be consistent with previously published data on human Nav channels. We then compared this data to the potency of block of Nav current measured in pyramidal neurons from neocortical brain slices. Interestingly, we found that it requires nearly 10-fold greater concentration of TTX over the IC_50_ to induce significant block of action potentials using a current-step protocol. In contrast, concentrations near the IC_50_ resulted in a significant reduction in AP firing and increase in rheobase using a ramp protocol. Surprisingly, a 20% reduction in action potential generation observed with 3 nM TTX resulted in significant block of seizure-like activity in the 0 Mg^2+^ model of epilepsy. Additionally, we found that approximately 50% block in pyramidal cell intrinsic excitability is sufficient to completely block all seizure-like events. These data serve as a critical starting point in understanding how fractional block of Nav channels affect intrinsic neuronal excitability and seizure-like activity. It further suggests that seizures can be controlled without significantly compromising intrinsic neuronal activity and determines the required fold over IC_50_ for novel and clinically relevant Nav channel blockers to produce efficacy and limit side effects.

## 1. Introduction

Voltage-gated ion channels form the basis for electrical activity in cells (Armstrong and Hille, 1998, Catterall, 1995). Specifically, voltage-gated sodium (Nav) channels are critical for action potential generation in excitable tissues (de Lera Ruiz and Kraus, 2015, Kwong and Carr, 2015). There are three predominant Nav channel genes expressed in the adult brain, denoted *SCN1A, SCN2A* and *SCN8A*, which encode the channels Nav1.1, Nav1.2 and Nav1.6, respectively (Wang et al., 2017, Goldin et al., 2000, Trimmer and Rhodes, 2004). These channels are expressed differentially in the central nervous system. Nav1.2 and Nav1.6 are expressed in excitatory neurons (Caldwell et al., 2000, Tian et al., 2014, Hu et al., 2009, Katz et al., 2018) while Nav1.1 is the major channel in inhibitory interneurons (Ogiwara et al., 2007, Catterall et al., 2010, Yu et al., 2006). These channels are critical for normal physiological function of neuronal network activity and alterations in their function can result in aberrant network activity. For example, gain of function mutations of both the Nav1.2 (Misra et al., 2008, Kearney et al., 2001, Liao et al., 2010, Ogiwara et al., 2009) and Nav1.6 (O’Brien and Meisler, 2013, Gardella and Moller, 2019, Johannesen et al., 2019) have been demonstrated to cause severe epilepsies while loss of function mutations of Nav1.1 is the main cause of Dravet syndrome (Catterall et al., 2010, Bender et al., 2012, Catterall, 2018, Cheah et al., 2012). Additionally, loss of function of Nav1.2 or Nav1.6 can result in neuronal disorders such as autism spectrum disorder, demonstrating that balance between excitation and inhibition must be maintained within the brain (Ben-Shalom et al., 2017, Larsen et al., 2015).

Despite the risk of causing cardiac, behavioral, or cognitive side effects, a large number of Nav channel blockers are used as anti-epileptic drugs (AEDs) (Zuliani et al., 2009, Khateb et al., 2021). All the currently available AEDs on the market that inhibit Nav channels are not selective between the Nav isoforms (pan-Nav). Our group has recently demonstrated the utility in animal models of a subtype specific Nav1.6 channel inhibitor which is effective at limiting seizures with a wider therapeutic margin compared with conventional pan-Nav AEDs (Johnson et al., 2022). However, these isoform selective Nav inhibitors are not yet available in the clinic. To further our understanding of Nav targeting AEDs, we sought to determine the minimal fractional block of Nav current required to limit seizure activity and how that impacts single cell intrinsic excitability. Here, we investigated this using the tool compound tetrodotoxin (TTX), which is approximately equipotent on all three predominantly expressed Nav channels in the adult brain and shows only mild state dependence which facilitates accurate titration of Nav channel availability across a brain slice without significant variation from diversity of membrane potential (Boccaccio et al., 1999). As this study would be conducted using mouse brain slices, we first characterized the potency of TTX on mouse Nav1.1, Nav1.2, and Nav1.6 channels expressed in HEK 293 cells using automated-patch clamp electrophysiology. We found the potency to be similar across Nav1.1, 1.2 and 1.6, consistent with the reported values on human sodium channels. Next, using brain slice electrophysiology, we also determined the efficacy of TTX in suppressing intrinsic neuronal excitability and showed that it required nearly 80% block of Na^+^ current to significantly reduce action potential firing in pyramidal cells and 100% block of Na^+^ current to fully block firing upon depolarizing current injections. However, using a ramp protocol, we found that approximately a 20% reduction in available sodium current can reduce action potential firing and right shift the rheobase. We next sought to determine the fractional block of Nav channels required to limit *ex vivo* seizure activity. Interestingly, we found that the zero Mg^2+^ model of epilepsy is more sensitive to sodium channel block than cell intrinsic properties, where a 20% reduction on Nav channels significantly reduced *ex vivo* seizure generation. These data demonstrate a discrepancy between somatic action potential firing and initiation of seizure-like activity. It also indicates that seizure control might be achieved without significantly compromising cell intrinsic properties and demonstrates a non-linearity in the relationship between Nav channel availability, neuronal excitability, and seizure-like activity suppression.

## 2. Material and Methods

### 2.1 Electrophysiology using HEK 293 cells

The potency of TTX on mouse Nav channel subtypes was assessed using a Qube 384 (Sophion) automated voltage-clamp platform. HEK-293 cell lines stably expressing Nav1.x channels correspond to the following GenBank accession numbers: mouse Nav1.1 (NM_018733.2), mouse Nav1.2 (NP_001092768.1) and mouse Nav1.6 (NM_001077499). The Nav β1 subunit (NM_199037) was co-expressed in all cell lines. Mouse Nav1.6 channels were also co-expressed with FHF2B (NM_033642) to increase functional expression. To measure the TTX-inhibition of Nav channels, the membrane potential was maintained at a voltage (−120 mV) where channels are mainly at rest followed by a depolarization for 10 sec to the empirically determined V_0.5_ for that cell followed by a test pulse to -20 mV for 5 ms to activate the Nav channels and quantify the TTX inhibition of the current. The protocol was repeated every 30 seconds. The V_0.5_ of steady state inactivation was determined using the Qube adaptive V_0.5_ function with a series of 500 ms depolarizing steps from a holding potential of -120 mV to -20 mV in 10 ms steps with a test pulse to -20 mV to evaluate voltage dependence of availability. The availability curve was fit with a Boltzmann charge-voltage equation to find the V_0.5_. To construct the TTX concentration response curves, baseline currents were established in vehicle for 5 minutes (0.5% DMSO) before addition of the appropriate TTX concentration for 5 minutes. Full inhibition response amplitudes were determined by adding tetrodotoxin (TTX, 300 nM) to each well at the end of the experiment. The fractional inhibition by TTX was calculated by normalizing the response in different concentrations of TTX to the baseline vehicle current and the full inhibition response in a supramaximal concentration of 300 nM TTX. Appropriate filters for minimum seal resistance were applied (typically >500 MΩ membrane resistance), and series resistance was compensated at 100%. Currents were sampled at 25 kHz and low pass filtered at 5 kHz.

### 2.2 Automated patch-clamp recording solutions

The recording solutions for Nav1.1, Nav1.2, and Nav1.6 cell line studies contained: Intracellular solution (ICS): 5 mM NaCl, 10 mM CsCl, 120 mM CsF, 0.1 mM CaCl_2_, 2 mM MgCl_2_, 10 mM HEPES (4-(2-hydroxyethyl)–1-piperazineethanesulfonic acid buffer), 10 mM EGTA (ethylene glycol tetra acetic acid); adjusted to pH 7.2 with CsOH. Extracellular solution (ECS): 140 mM NaCl, 5 mM KCl, 2 mM CaCl_2_, 1 mM MgCl_2_, 10 mM HEPES; adjusted to pH 7.4 with NaOH. Osmolarity for the ICS and ECS solutions was adjusted with glucose to 300 mOsm/kg and 310 mOsm/kg, respectively.

### 2.3 Animals

All animal handling and experimentation involving animals were conducted using approved protocols according to the guidelines of the Canadian Council on Animal Care (CCAC) and approved by the Xenon Animal Care Committee (XACC). Brain slice electrophysiology and MEA experiments were performed on male or female CF-1 ™ mice (3-5 weeks old, Charles River). All the mice colonies were maintained in the Animal Resource Facility at Xenon Pharmaceuticals. Mice were given food and water ad libitum and kept on a 12-h light/dark cycle.

### 2.4 Acute brain slice preparation

Mice were deeply anesthetized with isoflurane [5% (vol/vol) in oxygen] and decapitated. The brain was removed and transferred immediately into an ice cold oxygenated (95% O_2_, 5% CO_2_) sucrose cutting solution containing the following (in mM: 214 Sucrose, 26 NaHCO_3_, 1.6 NaH_2_PO_4_, 11 Glucose, 2.5 KCl, 0.5 CaCl_2_, 6 MgCl_2_, adjusted to pH 7.4. 300 μm-thick neocortical coronal/horizontal slices were collected for the patch-clamp studies and 350 μm-thick neocortical horizontal slices were collected for the MEA studies, cut using a VT1200 vibratome (Leica). The slices were then incubated in artificial cerebral spinal fluid (aCSF) containing (in mM): 126 NaCl, 3.5 KCl, 26 NaHCO_3_, 1.26 NaH_2_PO_4_, 10 Glucose, 1 MgCl_2_, 2 CaCl_2_, pH 7.4 with 95% O2-5% CO_2_ at 34 °C for 45 min prior to recording.

### 2.5 Brain slice patch-clamp electrophysiology

After incubation in aCSF for ∼ 45 min, individual slices were transferred to the recording chamber and constantly perfused at 2 ml/min with oxygenated aCSF maintained at 34°C. The neurons were visualized with infrared differential interference contrast (IR-DIC; Slicescope Pro 2000, Scientifica, UK) in combination with a 40X water immersion objective. Cortical pyramidal neurons were identified based on their characteristic large, triangular soma and single apical dendrite. Whole-cell patch clamp recordings were performed using a Multiclamp 700 B amplifier and signals were digitized and acquired using Digidata 1550B and pClamp 11 software (Molecular Devices). The recording chamber was grounded with an Ag/AgCl pellet. Patch pipettes (4-6 MΩ) were pulled from borosilicate glass using a P-70 micropipette puller (Sutter Instruments).

Intrinsic neuronal excitability was recorded in current-clamp mode using an internal recording solution containing the following (in mM): 120 K^+^ Gluconate, 10 HEPES, 1 MgCl_2_, 1 CaCl_2_, 11 KCl, 11 EGTA, 4 MgATP, 0.5 Na_2_GTP, pH adjusted to 7.2 using KOH and osmolarity adjusted to 290 mOsm/kg using D-mannitol. To record the intrinsic excitability of pyramidal neurons, the following synaptic blockers were added; NMDA receptor antagonist D-2-amino-5-phosphonovalerate (D-APV, 50 μM), AMPA receptor antagonist 6,7-dinitroquinoxaline-2,3-dione (NBQX, 10 μM) and GABA_A_ receptor antagonist gabazine (10 μM) were added to the aCSF. The liquid junction potential was +13.3 mV, and the data was reported without subtraction. Bridge balance values were monitored during recordings and cells displaying bridge balance values >20 MΩ were excluded from the analysis. Changes in intrinsic excitability were evaluated by using a current step and a ramp protocol. In case of step protocol, action potentials were evoked with 1 s long square-pulse current injections from -100 to 440 pA in 20 pA increments. A ramp protocol of 120 pA/ s (0 to 480 pA) was also used to evoke AP firing. Data under current clamp conditions were sampled at 50 kHz and low pass filtered at 10 kHz. Once baseline AP firing was determined in the extracellular recording solution (aCSF), the TTX with different concentrations was washed onto the slice for 15 min and the AP-generating protocol was repeated.

For recording Na^+^ currents in pyramidal neurons in voltage-clamp mode, the aCSF contained the following (in mM): 50 NaCl, 90 TEA-Cl, 10 HEPES, 2 CaCl_2_, 2 MgCl_2_, 3.5 KCl, 3 CsCl, 0.2 CdCl_2_, 4 4-aminopyridine, 25 Glucose, pH adjusted to 7.2 using NaOH. The internal solution contained the following (in mM): 110 CsF, 10 HEPES, 11 EGTA, 2 MgCl_2_, 0.5 NaGTP, 2 Na_2_ATP, pH adjusted to 7.2 with CsOH. Data under voltage clamp conditions were sampled at 20 kHz and low pass filtered at 2 kHz. Leakage and capacitive currents were automatically subtracted using a pre-pulse protocol (−P/4).

### 2.6 Multi-electrode array recordings

Recordings were performed on the 3Brain BioCAM DupleX system (Switzerland) using the 3Brain Accura HD-MEA chips with 4,096 electrodes at a pitch of 60μm. Slices were placed onto the electrodes with a harp placed on top to keep the slice pressed down gently to the recording electrodes. Slices were perfused continuously with a zero Mg^2+^ artificial cerebrospinal fluid (aCSF) to induce epileptiform activity. TTX was bath applied 10 minutes prior to washout of Mg^2+^ ions. Recordings were obtained from the entire neocortical slice. Recordings were performed at 33–36°C. The solutions were perfused at the rate of 5.0 mL/min. Signals were sampled at 10 kHz with a high-pass filter at 2 Hz. All analysis was done in our custom-made GUI (Mahadevan, 2022).

### 2.7 Data analysis and statistics

Brain slice patch clamp electrophysiology recordings were analyzed using Clampfit 11 (Molecular devices) and a custom-made GUI written in Python version 3.7. Python script can be made available upon request. The number of APs was measured and plotted as a function of current stimulus to allow comparison between vehicle and TTX treatment. Rheobase was measured as the minimum intensity of 1s current pulse required for initiation of AP. The AP threshold voltage was defined as the voltage at which the first derivative of the AP waveform (dv/dt) reached 10 mV/ms. The threshold was calculated at rheobase current. For AP waveform analysis, amplitude was measured from threshold to peak, and AP width measurements were made at 50% of the peak amplitude. AP upstroke velocity was assessed through phase plots that were generated using the membrane potential trace and the dV/dt trace.

All statistical analyses were performed using GraphPad Prism 7 software and differences with p-value < 0.05 were considered statistically significant. All the figures were created using diagrams, Inkscape 1.1. and all the data are expressed as mean ± SEM. The n value represents the number of cells tested for the patch clamp data or number of slices for the MEA data. For the statistical comparisons of the i-o curves, the difference in the cumulative area under the curves (AUC) for vehicle versus TTX treated conditions was evaluated by an unpaired two-tail test. The statistical analysis of the effect of TTX on the AP properties was evaluated using a One-way ANOVA (Figure 3 D-G) and paired two-tail test (Figure 4 C-F). MEA data was analyzed using a One-way ANOVA with a Dunnett’s test.

### 2.8 Pharmacological agents

D-AP5, NBQX and Gabazine were purchased from Hellobio. Tetrodotoxin (TTX) was purchased from Biotrend. All drugs were prepared as stock solutions and stored at − 20 °C; aliquots were thawed and added to aCSF to the working concentrations. Drugs were applied by bath perfusion at a flow rate of 2 ml/min.

## 3. Results

### 3.1 TTX inhibition of mouse Nav channels expressed in CNS

Previous studies reported the potency of TTX on rat or human Nav channel isoforms (Rosker et al., 2007, Tsukamoto et al., 2017, Zhang et al., 2013), however, the potency on the mouse neuronal Nav channel isoforms, to our knowledge, has not been reported in the same publication under the same experimental conditions. Here, using automated patch-clamp electrophysiology, we assessed the potency of TTX on the mouse Nav isoforms that are highly expressed in the CNS including: Nav1.1, 1.2, and 1.6 (Ogiwara et al., 2007, Tian et al., 2014, Dutton et al., 2013, Hu et al., 2009).

To assess the potency without biasing towards either the inactivated or resting states, we used a voltage-clamp protocol that holds channels at the V_0.5_ of inactivation, where the population of channels are approximately equally distributed between resting and inactivated states as described in the methods section. The V_0.5_ for each channel was as follows: mNav1.1 -54.4 ± 0.6 mV (n = 79), mNav1.2 -61.3 ± 0.9 mV (n = 45) and mNav1.6 -61.9 ± 0.8 mV (n = 42). This is to better simulate membrane potentials that are found in neurons that will typically have resting membrane potentials where channels are in mixed states. Representative current traces at approximately IC_50_ concentrations for each subtype tested are shown in Fig.1A (red trace). Complete block of Na^+^ current was observed by application of 300 nM TTX (green trace). The concentration-response relationship for the mNav1.1, 1.2 and 1.6 channels are plotted in Fig.1B. To construct the concentration-response, individual cells were exposed to single concentrations of TTX, normalizing the inhibition of each cell to its own vehicle baseline at each concentration. This data was then pooled, and the mean data was fit with a Hill-Langmuir equation. TTX potently inhibited mNav1.6 channels with an IC_50_ of 0.6 nM (95% CI: 0.5 to 0.7 nM). Inhibition of other mNav1.X isoforms required somewhat higher concentrations of TTX with IC_50_s of 2.6 nM (95% CI: 1.9 to 3.5 nM) for mNav1.1, and 7 nM (95% CI: 4.3 to 11 nM) for mNav1.2. The potency in mouse Nav channels was similar to those seen in human orthologues with low nanomolar IC_50_ of TTX on hNav1.6 and 1.1, and slightly higher potency on hNav1.2. The observed IC_50_s for TTX inhibition were 1.43 nM (95% CI: 0.8 to 2.3 nM) for hNav1.1, 2.5 nM (95%CI: 1.6 to 4 nM) for hNav1.2 and 0.5 nM (95%CI: 0.4 to 0.6 nM) for hNav1.6. These data on human Nav isoforms are consistent with those reported previously (Tsukamoto et al., 2017).

**Figure 1:**
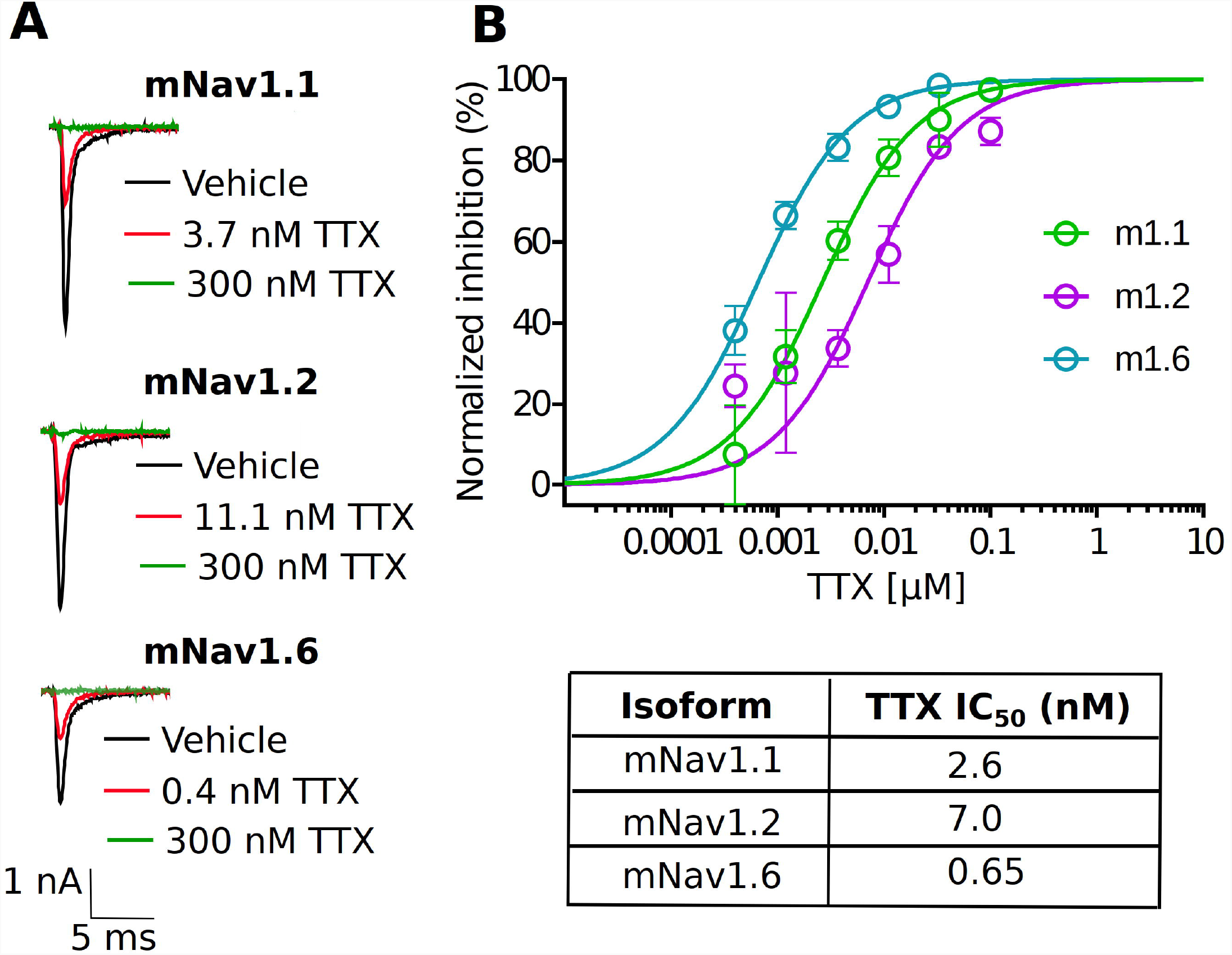
Potency of TTX inhibition on the mouse neuronal Nav channels. (***A***) Representative traces of voltage-clamp recordings of vehicle (black) and the concentration close to the TTX IC_50_ (red) for mouse Nav channel isoforms (mNav1.1, mNav1.2 and mNav1.6) heterologously expressed in HEK293 cells. For all the tested Nav channel isoforms, 300 nM TTX (green trace) showed complete block of Na^+^-current. (***B***) Concentration-response relationship for TTX inhibition of mouse Nav1.1, Nav1.2 and Nav1.6 channel isoforms. Each concentration represents the mean of n = 3-12 cells. The solid line is the best fit of the average data to the Hill equation yielding IC_50_ = 2.6 nM for mNav1.1, 7.04 nM for mNav1.2 and 0.65 nM for mNav1.6.

### 3.2 Inhibition of TTX on Na_+_ currents and AP firing in pyramidal neurons in brain slices

After the characterization of TTX potency in heterologously expressed mouse Nav channels, we moved to electrophysiological analysis of neuronal Nav currents in the physiologically appropriate environment of acute brain slices. We made whole-cell patch clamp recordings of excitatory pyramidal neurons from coronal slices, in which Nav1.2 and Nav1.6 channels are predominantly expressed (Tian et al., 2014, Hu et al., 2009, Katz et al., 2018, Spratt et al., 2021). Different concentrations of TTX (3, 10 and 30 nM) were tested to measure the sensitivity of Na^+^ currents to TTX. Representative whole-cell Na^+^ currents recorded from pyramidal neurons in response to a typical voltage-clamp step protocol is shown in Fig. 2A. Application of increasing concentrations of TTX resulted in concentration-dependent inhibition in the Na^+^ peak current. Na^+^ currents were reduced to 83.05 ± 5.3% of vehicle in 3 nM TTX (n = 6), 56.7 ± 6.6% of vehicle (n = 6) in 10 nM TTX; and 19.1 ± 6.2% of vehicle (n = 5) in 30 nM TTX (Fig.2A and B). The concentration-response curve for TTX on Na^+^ currents from pyramidal neurons was constructed and fitted with a Hill-Langmuir equation giving an IC_50_ value of 9.6 nM (95% CI: 7.5 to 12 nM) (Fig.2B). This result is comparable to the potency of TTX on heterologously expressed channels (Fig. 1) and other reported effects of TTX on Na^+^ currents in isolated CA1 pyramidal neurons (IC_50_ = 6.4 nM) (Madeja, 2000) and GABAergic neurons (IC_50_ = 7.2 nM) of the substantia Nigra obtained in nucleated patches (Seutin and Engel, 2010).

**Figure 2:**
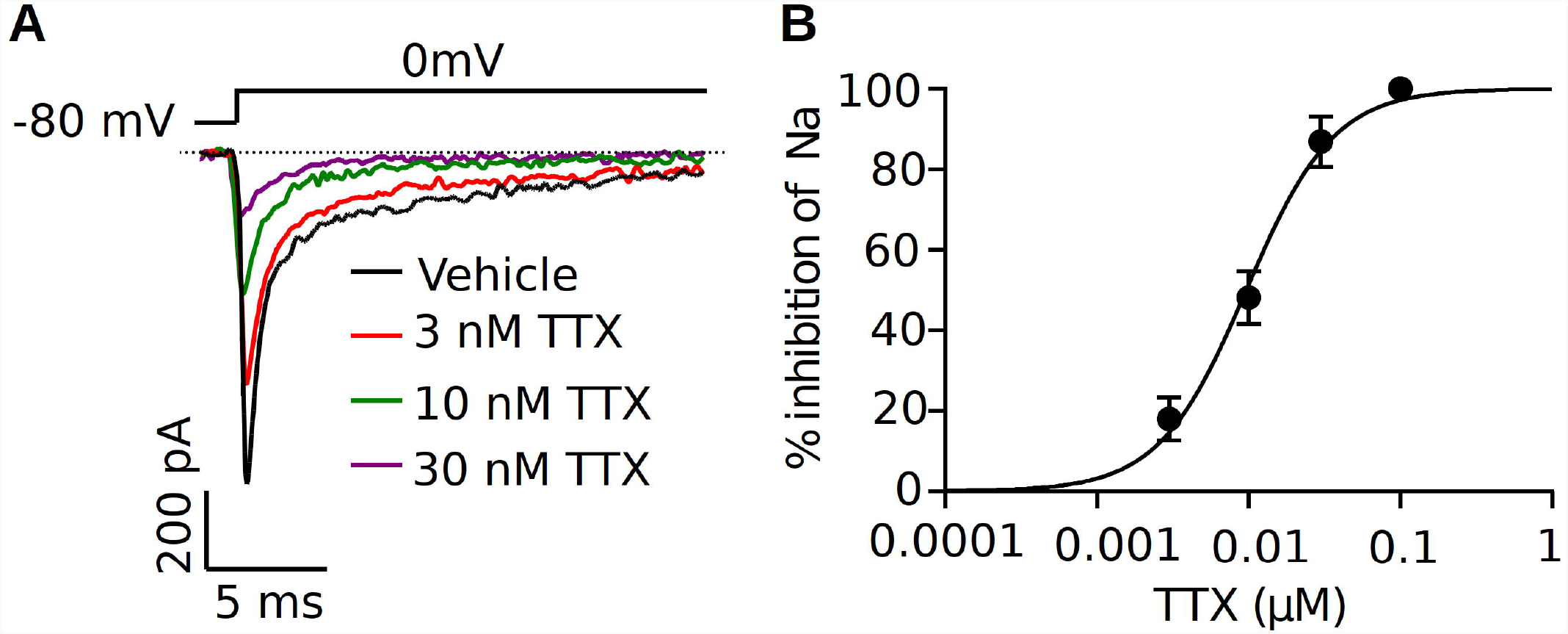
Increasing concentrations of TTX resulted in a concentration-dependent inhibition of Na^+^ currents in mouse cortical pyramidal neurons. **(*A*)** Representative traces of Na^+^ currents from pyramidal neurons recorded in vehicle condition and in presence of TTX at concentrations of 3, 10 and 30 nM. Inhibition by TTX of Na^+^ currents were elicited by a 20-ms pulse depolarized from a holding potential of -80 mV to 0 mV. (***B*)** Concentration-response relationship for TTX inhibition of Na^+^ current in pyramidal neurons. Each point shows Na^+^ peak current relative to vehicle averaged over 6 neurons, except for 100 nM TTX (complete block in 1 cell). The solid line is the best fit of the average data to the Hill equation yielding IC_50_ = 9.6 nM.

To determine the potency of TTX on AP firing, we next performed current-clamp recordings from the soma of pyramidal neurons in the presence of the synaptic blockers NBQX (10 μM), AP5 (50 μM) and Gabazine (10 μM). Fig.3A shows the representative voltage responses to 220 pA current injection step in vehicle conditions and in the presence of different concentrations of TTX (3, 10 and 30 nM). Application of 10 nM TTX significantly reduced the number of APs by almost 60% across a range of stimuli and in 30 nM TTX, depolarizing current did not elicit APs (Fig 3B). Interestingly, the lowest concentration of TTX (3 nM) blocked ∼ 20% of the Na^+^ current (Fig.2) but showed little or no effect on the AP firing (Fig.3B). A concentration-response relation for TTX block on AP firing at maximal current stimulus (440 pA) revealed the half-maximal blocking concentration to be 7.9 nM (95% CI: 5.0 to 11 nM) (Fig. 3C).

**Figure 3:**
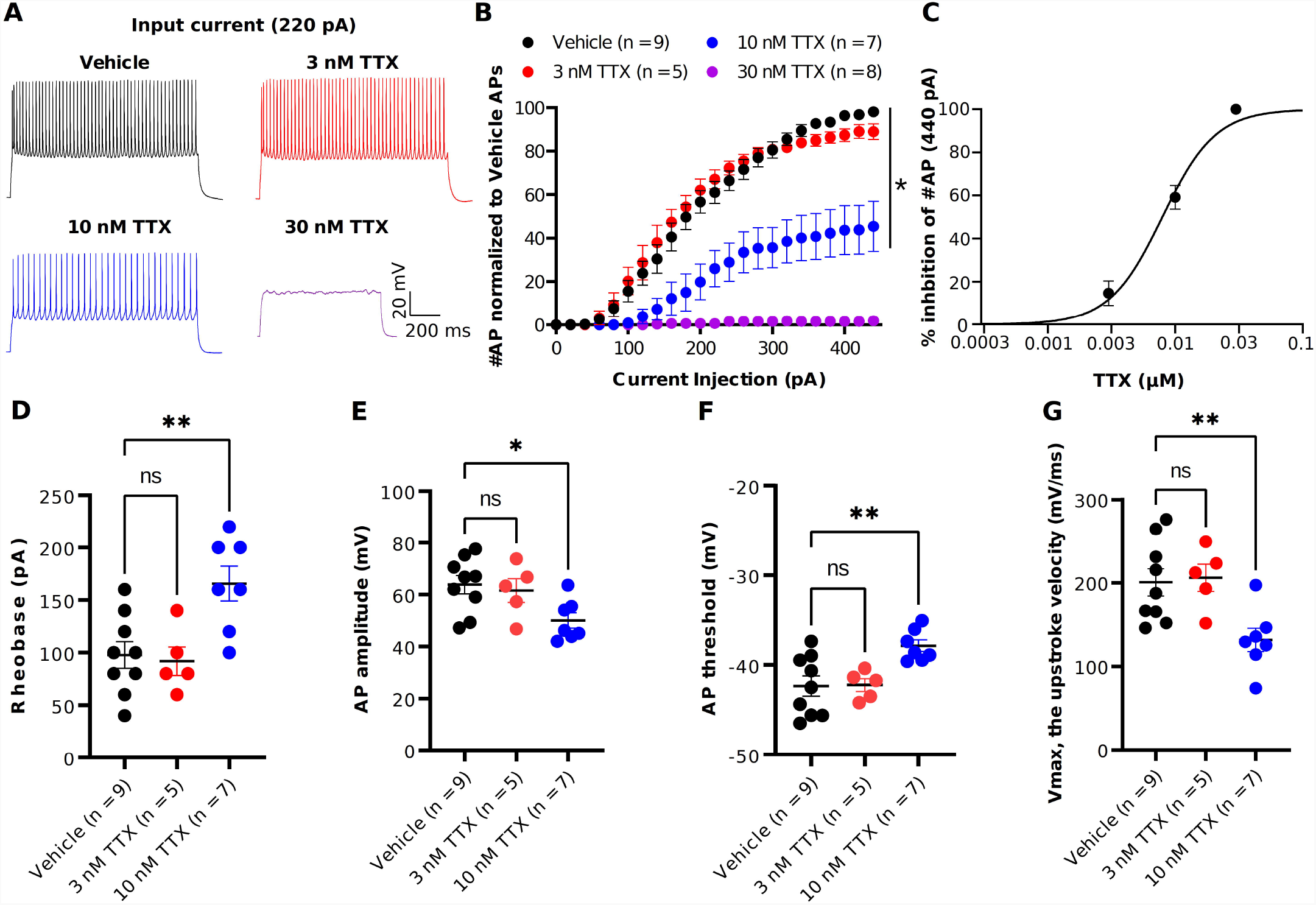
TTX suppression of intrinsic excitability in cortical pyramidal neurons. (***A***) Representative voltage traces recorded from a cortical pyramidal neuron at a current injection of 220 pA in vehicle (black) and in presence of TTX at concentraions of 3 (red), 10 (blue) and 30 nM (magenta). (***B***) Mean data of number of APs as a function of various current injection for pyramidal neurons with and without TTX application. The area under the curve (AUC) for current injections > 220 pA was significantly decreased for 10 and 30 nM TTX. The AUC for vehicle versus TTX treated conditions was evaluated by an unpaired two-tailed t-test (\**p*<0.005). (***C***) Concentration-response relationship for TTX inhibition of AP firing at a current injection of 440 pA. Each concentration represents the mean of n = 5-9 neurons. The solid line is the best fit of the average data to the Hill equation yielding IC_50_ = 7.9 nM. (***D-G***) Scatter plots showing the AP properties of pyramidal neurons include rheobase (**D**) AP amplitude (**E**), AP threshold (**F**) and upstroke velocity (**G**). 10 nM TTX significantly increased rheobase and AP threshold whereas AP amplitude and upstroke velocity was significantly decreased (one-way ANOVA, \**p*<0.005).

We next examined the passive and active parameters of AP firing. The resting membrane potential and input resistance were not altered by application of TTX (data not shown). Application of 10 nM TTX increased significantly the rheobase (vehicle: 112 ± 25.7 pA, n = 9; 10 nM TTX: 170 ± 15.1 pA, n = 7, p <0.05, Fig.3D) and AP threshold (vehicle: -41.4 ± 1.3 mV, n = 9; 10 nM TTX: - 36.1 ± 1.7 mV, n = 7, p <0.05, Fig.3F) whereas the AP amplitude (vehicle: 63.0 ± 3.2 mV, n = 9; 10 nM TTX: 48.1 ± 3.1 mV, n = 7, p <0.05, Fig.3E) and maximum AP velocity (vehicle: 200.6 ± 16.2 mV/ms, n = 9; 10 nM TTX: 132.4 ± 12.1 mV/ms, n = 7, p <0.05, Fig.3G) were significantly decreased. Together, these data suggest that it required nearly 80% block of Na^+^ current to significantly reduce AP firing and properties. Interestingly, low concentration of TTX showed very little effect of AP firing, whereas we did observe a significant block of Na^+^ current at 3 nM TTX (Fig.2).

Previous studies in Purkinje neurons (Raman and Bean, 1999, Swensen and Bean, 2005) reported that low concentrations of TTX (∼ 3 nM) significantly impacted AP firing. As shown in Fig.3, when measured using square-step current injection protocol, 3 nM TTX did not affect the number of APs or AP firing properties of pyramidal neurons. To further investigate the effects of TTX on neuronal excitability, we recorded AP firing using a slow ramp current injection. Fig.4A shows the representative voltage responses to a current ramp (4 s ramp from 0 to 480 pA; 120 pA/s) recorded from a pyramidal neuron in vehicle condition (black trace) and in the presence of 3 nM TTX (red trace). Unlike the step protocol (Fig.3), we found significant reduction in the number of APs with addition of 3 nM TTX compared to that of vehicle (Fig.4B) suggesting the ramp is a more sensitive assay of Nav channel contributions to excitability. In addition, the rheobase and latency to first spike was increased significantly over vehicle whereas the AP amplitude was decreased. (Fig.3C, 3D, 3E). This result suggests that 20% reduction of Na^+^ current is sufficient to influence neuronal excitability of pyramidal neurons when measured using a ramp protocol.

**Figure 4:**
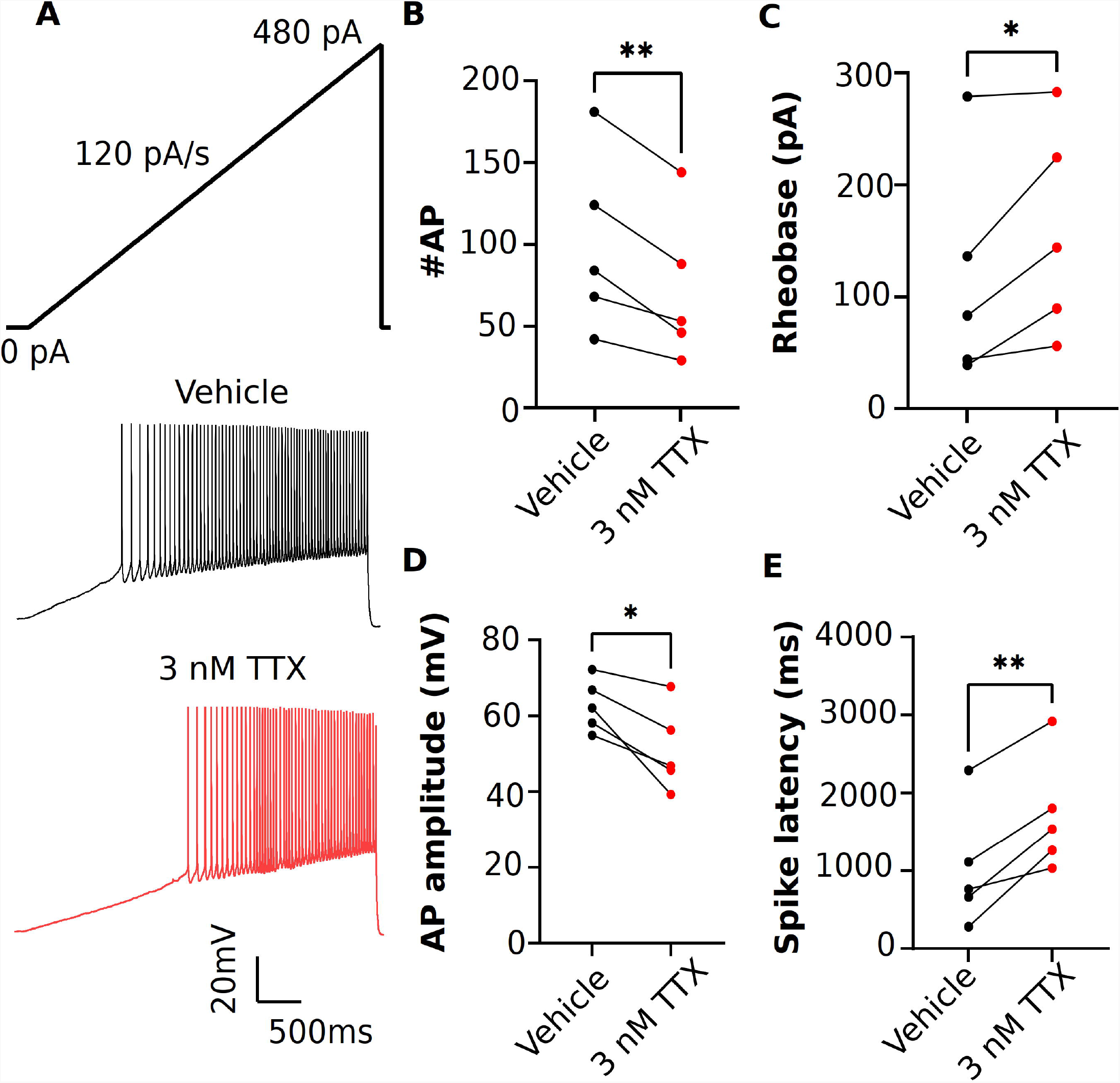
Effects of low concentration of TTX on intrinsic excitability of pyramidal neurons using a current ramp injection. (***A***) Representative voltage traces of pyramidal neurons recorded in response to a current injection ramp (4 s ramp from 0 pA to 480 pA; 120 pA/s) without (black) and with (red) application of 3 nM TTX. (***B***) Scatter plot showing significant impact on the number of APs with application of 3 nM TTX. (***C-E***) Using this protcol, we found that the rheobase (C), AP amplitude (D) and spike latency (E) was also significantly altered with application of 3 nM TTX (paired 2 tailed t-test, *p< 0.005; n = 5 neurons).

### 3.3 Inhibition of seizure-like events in TTX shows a clear concentration-dependent response

We next assessed the potency of TTX block of *ex vivo* seizure-like events, using a zero-magnesium aCSF solution. Removal of Mg^2+^ ions typically results in seizure-like events in ∼10-20 minutes. A representation of the seizure-like activity and the effects of various concentration of TTX is shown in Fig. 5. Fig. 5B displays representative raster plots from a slice in vehicle, and then 3 and 10 nM TTX respectively, demonstrating a decrease in epileptiform activity with increasing TTX concentration. The time-series traces over the entire 40-minute recording period further demonstrate the loss of seizure-like events between vehicle and TTX bathed slices, suggesting 10 nM TTX is sufficient to block all seizure-like events (Fig. 5C). Taking a closer look at the first seizure-like event from each treatment group, we also found that increasing TTX concentration appears to reduce higher frequency components during the seizure-like event, suggesting less neuronal recruitment during these pathological events (Fig. 5D, 5E).

**Figure 5.**
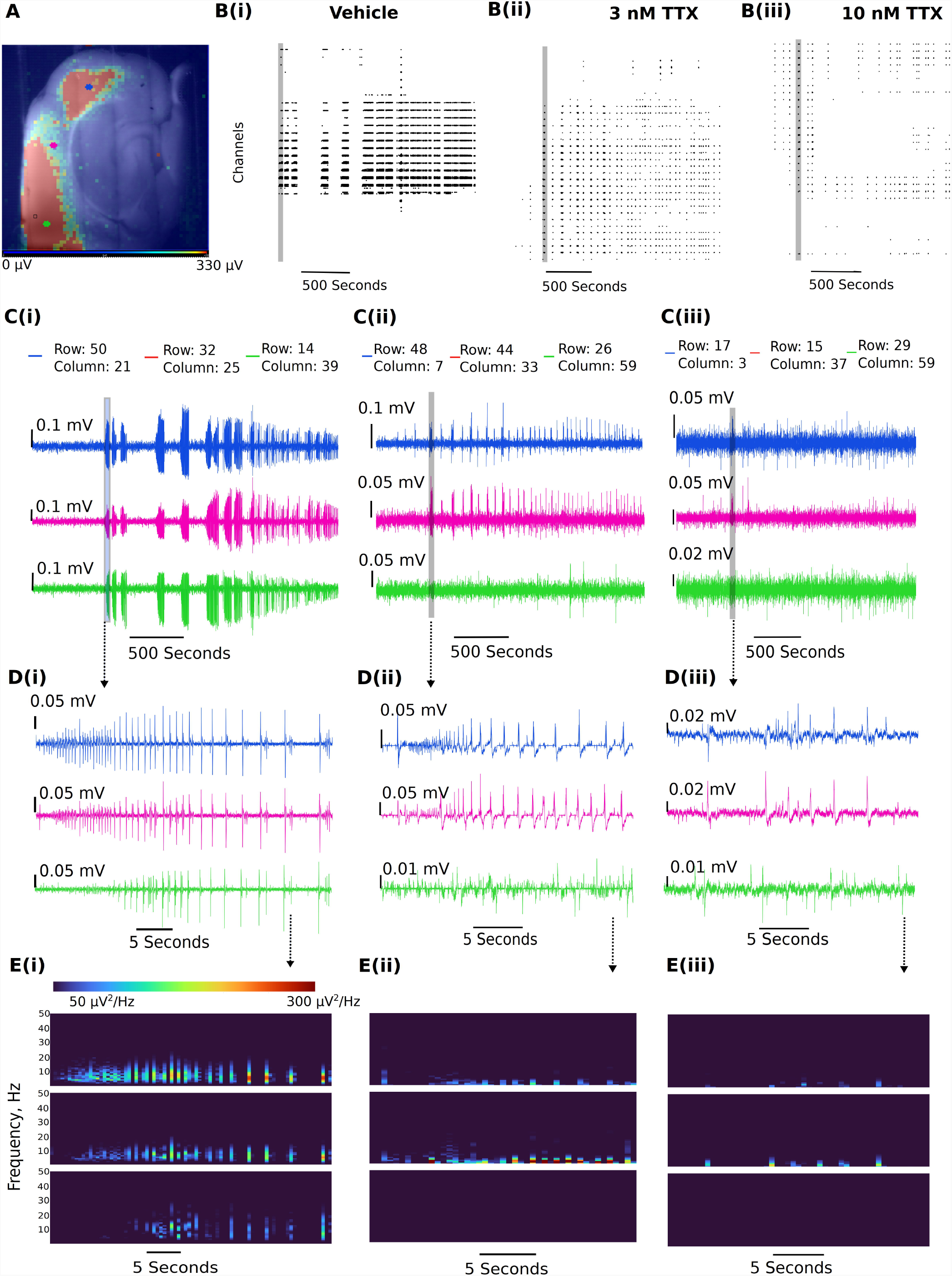
Representative reduction in seizure-like events following sodium current block by TTX. A) Example slice from a multi-electrode array recording during a seizure-like event. Colored dots (blue, pink, and green dots) represent the approximate location of the selected traces. B) Example raster plots from the entire recordings for the (Bi) 0 Mg^2+^ vehicle, the (Bii) 3 nM TTX, and the (Biii) 10 nM TTX. C) Example traces taken from either end of the neocortical slice (blue and green traces) and a trace selected from the middle of the slice (pink trace) for the (Ci) 0 Mg^2+^ vehicle, the (Cii) 3 nM TTX, and the (Ciii) 10 nM TTX. D) Zoomed in view of the first seizure-like events from each of the three treatment groups. E) Spectral plots of the zoomed in seizure-like events in μV^2^.

3 nM TTX is sufficient to significantly block seizure-like events but 10 nM is required to completely prevent seizures from developing (Fig 6). These data demonstrate that only a 20% reduction in Na^+^ current in pyramidal cells can produce a significant reduction in seizure-like events, even when this results in a rather modest block of action potential firing in pyramidal cells (see Fig 3 and 4). However, to fully achieve block of seizure-like events, it appears to require a 50% reduction in pyramidal cell Na^+^ current. In summary, these data demonstrate that only a partial block of pyramidal cell firing is required to achieve significant reductions in seizure-like events.

**Figure 6.**
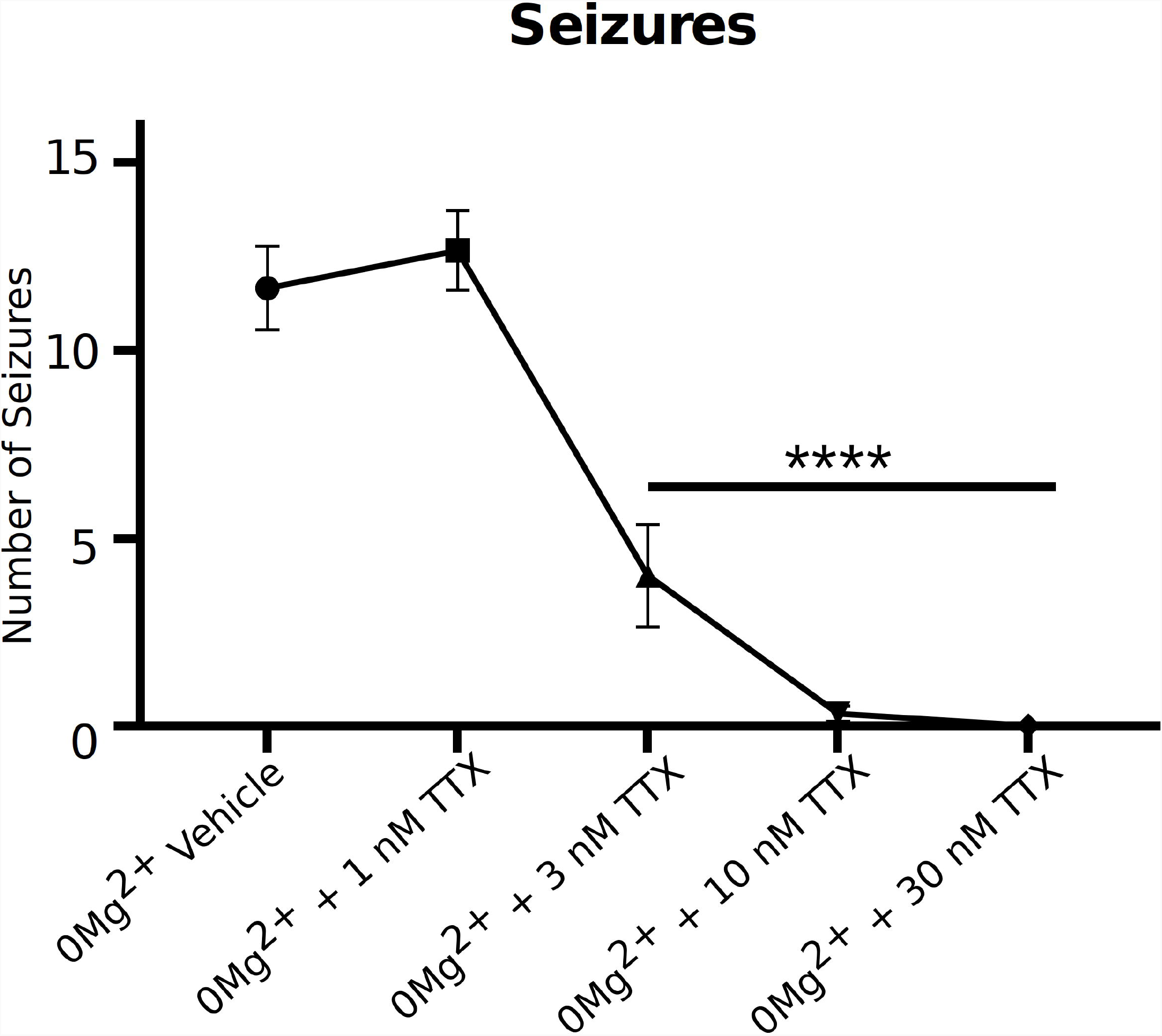
TTX reduces seizure-like events in a clear concentration dependent manner. A) Seizure-like events are reduced in a TTX concentration-dependent manner (one-way ANOVA with a Dunnett’s test, * = p<0.0001, n = 6).

## 4. Discussion

In the present study, we have demonstrated that relatively subtle reductions in pyramidal cell sodium current (∼20%) is sufficient to significantly reduce *ex vivo* seizure-like activity. Interestingly, this 20% reduction in sodium current has relatively little effect on intrinsic excitability of pyramidal cells, where we found blocking ∼ 80% of sodium current was required to impair neuronal firing with a standard step protocol. However, we also found that TTX sensitivity can be dependent on how the cell receives the current input, as a ramp protocol (Fig. 4) demonstrated that neuronal action potentials can be significantly impaired at lower concentrations of TTX than a step protocol. Overall, our data suggests that subtle impairment of neuronal action potential firing with sodium channel blockers might be sufficient to reduce seizures, a finding that can be important for exploration of novel anticonvulsant compounds and when determining a therapeutically relevant dosing of compounds. To our knowledge, the current study is the first to determine the fractional block of Nav channels required to reduce neuronal excitability at both the cellular and network level. In addition, we have also assayed the potency of TTX at different levels of network and neuronal activity, providing novel insights into how pore Nav channel blockers behave during different forms of stimulation.

Of the available Nav inhibitors, we believe that TTX has good properties for tuning neuronal Nav currents due to the relatively low state dependence of around 3X (Boccaccio et al., 1999) compared with Nav targeting AEDs, which show much greater state dependence of at least 10X (Kuo et al., 1997, Kuo and Bean, 1994). Potency is therefore relatively stable across the ranges of membrane potentials that may be found between different neurons and sub compartments. Several studies have demonstrated the potency of TTX on human and rat Nav1.6, 1.2, and 1.1 channels (Rosker et al., 2007, Tsukamoto et al., 2017, Zhang et al., 2013), however, this is the first report (to our knowledge) demonstrating the potency of TTX on these three mouse channel isoforms in an exogenously expressed system. Our results on the mouse neuronal Nav channels are in line with what has been reported in both human and rat channels. These results allow for greater understanding and interpretation of data from mouse tissue when using TTX for partial block of Nav currents. It is also important to note that while we and others generally report TTX to be roughly equipotent on all three predominantly expressed Nav channels in the CNS, we did find an approximate 10X different in potency between inhibition of Nav1.6 vs Nav1.2 channels, consistent with the previous report on the inhibitory effects of TTX on human Nav 1.6 and Nav1.2 isoforms (Tsukamoto et al., 2017). Additionally, when assessing results from neuronal pan-sodium channel blockers, one must consider neuronal cell-type distribution of these channels. Nav1.6 and Nav1.2 channels are predominantly expressed in pyramidal cells (Caldwell et al., 2000, Tian et al., 2014, Hu et al., 2009, Katz et al., 2018) whist Nav1.1 channels are the dominant sodium channel isoform in interneurons, particularly in the so-called fast spiking parvalbumin positive interneurons (Ogiwara et al., 2007, Yu et al., 2006). This data would suggest that pan-Nav channel blockers should reduce network excitability of all neuronal cell types, however, the network effect from this can be more complicated. Potent block of GABAergic neurons via suppression of Nav1.1 channels could result in pro-epileptic effects, as has been previously demonstrated (Patel et al., 2020). Nevertheless, pan-Nav channel blockers are effective anticonvulsant drugs in many situations, but their efficacy may be limited in situations with already impaired interneuron function. This is most clearly demonstrated by the fact that clinical approved sodium channel blockers are contraindicated in treatment of seizures associated with Dravet syndrome (Brunklaus et al., 2012, Guerrini et al., 1998, Horn et al., 1986), a condition believed to be due to a reduction in Nav1.1 current. Interestingly, there is a body of literature that suggests in certain situations, interneurons can trigger seizure-like activity (Chang et al., 2018, Librizzi et al., 2017, Magloire et al., 2019, Shiri et al., 2015, Yekhlef et al., 2015). However, in the majority of physiological situations, it should be advantageous to preserve interneuron activity, where they provide on demand fast feed-forward inhibition to limit spreading seizures (Trevelyan et al., 2006, Cammarota et al., 2013, Parrish et al., 2018, Sessolo et al., 2015, Parrish et al., 2019). Interestingly, there is evidence that TTX has milder effects on somatic AP firing of fast-spiking interneurons and that other pan-Nav blockers have a limited effect on the GABAergic network compared to the Glutamatergic network (Kaneko et al., 2022, Pothmann et al., 2014). This could explain why we found that lower concentration of TTX is required to significantly limit seizure-like events compared to concentrations required to limit intrinsic excitability of pyramidal cells. Additionally, while our pyramidal cell intrinsic excitability study assays axonal initial segment (AIS) and somatic excitability, seizure like-events are network level events, where dendrites, axons, and postsynaptic release of neurotransmitters are also critical components of seizure propagation. For example, dendritic plateauing has been shown to be necessary for seizure generation, suggesting important network changes required for this type of pathological activity (Graham et al., 2021). Furthermore, axonal propagation of action potentials in neurons has been demonstrated to be more sensitive to sodium channel block than somatic action potentials (Kaneko et al., 2022, Hu and Jonas, 2014). Therefore, it is possible that lower concentrations of TTX impairs these other important network components required for seizure genesis more profoundly than it does at the cell soma and the AIS, but this has not been studied in detail.

In conclusion, we have characterized the fractional block of sodium channels required to reduce the neuronal excitability of pyramidal cells and *ex vivo* seizure-like events using the tool compound TTX. This data could be helpful in assaying dose requirements to reduce seizures with clinically relevant Nav channel inhibitors. Currently, only pan-Nav channel blockers are available for clinical use, which, like TTX, are not highly selective. Whilst we await the assessment of isoform selective Nav inhibitors, we need a better understanding of relevant exposures of pan-Nav blockers that may reduce seizures while limiting side effects. Our data suggests that it might be possible to use low levels of these pan-Nav blockers to reduce seizures without greatly impairing action potential firing in pyramidal neurons. However, it remains to be seen if other pan-Nav blockers will behave in a similar manner, with somewhat mild effects on cell intrinsic excitability whist reducing seizure activity. Nevertheless, this report demonstrates an important concentration response of seizure-like events to sodium channel inhibition and sets the groundwork for greater understanding of fractional block of compounds required for inhibition of pathological activity, whist trying to limit their impact on physiological properties of the network.

## Data availability statement

The raw data supporting the conclusions of this article will be made available by the authors, without undue reservation.

## Acknowledgments

We would like to thank the entire Xenon family for their support of this work.

## Author contributions statement

RP and SJG conceived this work. AM wrote the analysis code. ST, MW, SL, MS, and RP collected the data. ST, MW, SL, SJG, AM, and RP analyzed the data. ST, MW, and RP wrote the manuscript. All authors approved the final draft.

